# Haplotype synthesis analysis in public reference data reveals functional variants underlying known genome-wide associated susceptibility loci

**DOI:** 10.1101/027276

**Authors:** André Lacour, Tim Becker

## Abstract

The functional mechanisms underlying disease association identified by Genome-wide Association Studies remain unknown for susceptibility loci located outside gene coding regions. In addition to the regulation of gene expression, synthesis of effects from multiple surrounding functional variants has been suggested as an explanation of hard-to-interpret associations.

Here, we define filter criteria based on linkage disequilibrium measures and allele frequencies which reflect expected properties of synthesizing variant sets. For eligible candidate sets we search for those haplotypes that are highly correlated with the risk alleles of a genome-wide associated variant.

We applied our methods to 1,000 Genomes reference data and confirmed Crohn’s Disease and Type 2 Diabetes susceptibility loci. Of these, a proportion of 32% allowed explanation by three-variant-haplotypes carrying at least two functional variants, as compared to a proportion of 16% for random variants *(P* = 2.92 · 10^−6^). More importantly, we detected examples of known loci whose association can fully be explained by surrounding missense variants: three missense variants from MUC19 synthesize rs11564258 (L0C105369736/MUC19, intron; Crohn’s Disease). Next, rs2797685 (PER3, intron; Crohn’s Disease) is synthesized by a 57 kilobase haplotype defined by five missense variants from PER3 and three missense variants from UTS2. Finally, the association of rs7178572 (HMG20A, intron; Type 2 Diabetes) can be explained by the synthesis of eight haplotypes, each carrying at least one missense variant in either PEAK1, TBC1D2B, CHRNA5 or ADAMTS7.

In summary, application of our new methods highlights the potential of synthesis analysis to guide functional follow-up investigation of findings from association studies.

## 1. Introduction

Genome-wide Association Studies (GWAS) [1, 2] detected a multitude of genetic risk variants associated with complex diseases and phenotypes. For a great portion of these variants, the underlying biological mechanisms are still unknown. In public databases (GWAS Catalog, dbGene, etc.), the gene closest to the strongest association signal is provisionally listed as the susceptibility gene. However, other genes nearby might embody the true functional origin and cause the association signal via more or less complicated patterns of linkage disequilibrium (LD). As pointed out by the authors of [3, 4] there is no guarantee, that causal variants are mandatory in particular high LD with the top association signal (we will call it the *tag variant* throughout this work). For instance, interaction between multiple causal loci interfere with the ability to find either of them separately, but create a strong signal at a distantly linked marker [5].

Goldstein [6], see also [7], suggested that an association of a common variant with a complex disease can be synthetically created by multiple rarer functional variants from a surrounding genomic region. In this case the rarer variants occur more often or exclusively on a haplotype branch carrying a specific allele of the tag variant, generating in this way the strongest association signal at this locus. This situation is depicted in figure 1 of [8]. The idea can be quantified by checking whether the LD measures between tag variant and candidate variants yield |*D′|* ≈ 1 and *r*^2^ not large [6, 9]. Examples for such *synthetic associations* have been reported from GWAS [10, 11, 12, 13, 14] and sequencing studies [8]. A statistical method to test a given set of variants for synthetic association with quantitative trait has been described by [9].

The concept of synthetic association has also given rise to some debate: some authors considered the ubiquity of synthetic associations to be unlikely [15], while others discussed whether synthetic associations should have already been detected by linkage analysis [16, 17] and whether a rare-only model for synthetic association is applicable to a lot of GWAS findings [17, 18]. In addition, the expected properties of synthetic associations were empirically assessed via simulation studies [4, 19], with partially contradictory results. The authors of [4] stated, that rarer variants that contribute to a synthetic association might be as far as 2.5 Megabases (Mb) away, whereas the authors of [19] reported that in 90% of their simulations at least one rare causal variant was already captured within a window of size 100kilobases (kb). In any case, it can be stated, that until now there is no complete understanding of which role synthetic associations actually play for the etiology of complex diseases, how frequent the phenomenon of synthesis really occurs, and whether it is rather build up by a few low-frequency and common variants or a lot of rare variants.

In view of the lack consensus judgment on the relevance of synthetic association, we started an empirical evaluation of the frequency of the phenomenon. Until now, no methods have been provided to systematically search for sets of variants that synthesize the association of a common variant. A major reason for this is the computational load. Already within a ±100 kb region surrounding a susceptibility locus, typically 2,000 variants are to be expected. With *n* eligible variants in an identified trait-related susceptibility region, there are 2*^n^* – 1 variant sets to be investigated. Even when only sets with, for instance, up to 6 variants shall be tested for potential synthesis, 8.84 · 10^16^ different sets have to be investigated. In view of the large number, an efficient search engine is prerequisite for the detection of synthetic associations. Identification of such variant sets is highly relevant for the follow-up of association signals that were produced by GWAS or next generation sequencing (NGS) association studies, in order to come closer to the variants with disease relevant biological function [20].

This work is organized as follows: in section 2 we define filtering criteria under which potential variant sets for synthetic associations are selected, and we motivate the haplotype–tag variant correlation as a measure of synthesis. We apply our methods to data from the 1,000 Genomes Project [21] in section 3: a) frequency of the phenomenon of synthesis with broad-sense functional variants compared between random micro-array variants and known susceptibility loci of Crohn’s Disease and Type 2 Diabetes; b) identification and description of synthetic associations given by missense variants. A detailed discussion of our findings and conclusions are given in section 4.

## 2. Methods

We consider an LD region that is associated with a disease phenotype. The top association signal (tag variant) has been reported according to a GWAS catalog or a consensus meta-analysis. We assume that variant genotype data, either from public reference data, GWAS, imputed GWAS or NGS association studies, for the tag variant and a sufficiently large surrounding region are available. We advise to include variants in a window size between 2 and 5 Mb with the tag variant in its center.

### 2.1. Filtering rules

Let *a_i_, A_i_* be the alleles of variant *s_i_*. Let *f*(*a_i_*) be the allele frequency of *a_i_* and let *h*(*a_i_a_j_*) be the frequency of the 2-variants haplotype. The alleles *a_i_* and *a_j_* are said to be *in-phase* if D = *h*(*a_i_a_j_*) − *f* (*a_i_*) *f*(*a_j_*) > 0. From the data we calculate the allele frequencies and the LD measures *r*^2^ = *D*^2^/(*a_i_a_j_A_i_A_j_*) and *D′* = *D/D_max_,* where *D_max_* = min(*a_i_a_j_, A_i_A_j_*) if *D* > 0 and *D_max_* = min(*a_i_Aj, A_i_a_j_*) if *D* < 0. Note that *D*′ comprises a leading sign.

We employ the following allele frequency criteria for synthetic association. Let *S* = {*s*_1_,…, *s_k_*} be a set of *k* variants and let *s ∉ S* be the tag variant. Let *D′*(*a_i_, a_j_*), *r*^2^(*a_i_, a_j_*) be the pairwise LD measures for two variants *s_i_, s_j_*. Let 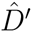 and 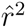 be predefined intervals, that quantify the conditions |D′| ≈ 1 and *r*^2^ not large [6, 9] and *t* ≳ 1 be a tolerance parameter.

Let *a* be the risk allele of *s*, i.e. the allele with an reported OR > 1. Let *a_i_* be the alleles of *s_i_* that are in-phase with *a. S* is said to be a **risk candidate set** if

1. 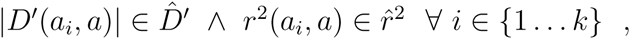
2. 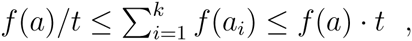
3. 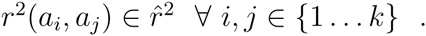

In the same way, we can regard *A* to be the protective allele of *s*, i.e. the allele with an reported OR < 1. Let now *A_i_* be the alleles of *s_i_* that are in-phase with *A*. *S* is then said to be a **protective candidate set** if the above criteria hold, whereupon the second condition is replaced by

2.) 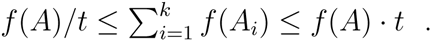

One may also see this twofold search from a different perspective: once we seek candidate sets that are synthesized by the minor allele of the tag variant, which means *D′*(*a_i_, a*) *>* 0, and another where the sets are synthesized by the major allele of the same, *D′*(*a_i_, a*) < 0.

In the context of this work, we prune variants for *r*^2^ = 1, while we do not remove variants that are marked to have known functional consequences. As parameters we choose *t* = 1.1 and filter for an LD space of 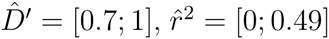. More details on our implementation are given in Appendix A.

### 2.2. Tag variant – set haplotype correlation

The goal of our approach is to find variant sets that explain a given tag variant via synthesis, in particular to find haplotypes that are in nearly perfect LD with one of the alleles of the tag variant. For a candidate set *S* of tag variant s we phase and reconstruct the haplotypes from the genotypes employing the EM-algorithm using maximum-likelihood estimation of haplotype frequencies according to [22, 23]. Here, we use an implementation that improves our previous implementation in FAMHAP [24]. We evaluate the haplotypes in a binary storing version, which is presented in detail in Appendix B. Haplotypes with a frequency below a cutoff will not be considered. From the reconstructed haplotypes with their estimated frequencies we compose dichotomized *haplotype markers* consisting each of one haplotype versus all others. Then we calculate the Pearson product-moment correlation coefficient of that allele *x* of tag variant *s* which is tested for being synthesized by *S* and each haplotype marker *h* of candidate set *S*

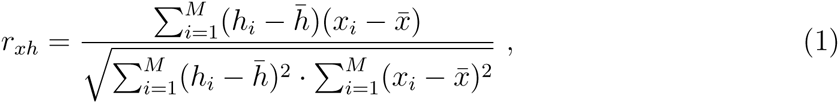

where *M* is the number of individuals, *x_i_* ∈ {0,1, 2} is the *i*th individual’s allele count of *s* and *h_i_ = h_i_*(*a_i_… a_k_*) ∈ [0; 2] is the frequency of the haplotype with set variant alleles *a_j_* for individual *i* from the maximum-likelihood estimation. 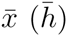 denote the mean values of all *x_i_* (*h_i_*). Synthesis is established if |*r_xh_*| ≈ 1. Note that *r_xh_* comprises a phase information in terms of a leading sign. In case of *k* = 1, *r_xh_* is equal to *r*, with *r*^2^ the standard pairwise LD measure.

## 3. Data analysis and results

### 3.1. Frequency of the synthesis phenomenon

Nearly perfect pairwise LD (*r*^2^ ≈ 1) between neighboring variants is a common phenomenon. Likewise, perfect LD between a single variant and a haplotype marker defined by a set of variants is likely to exist in regions of strong LD. In order to assess the frequency of the phenomenon, we randomly selected 1,000 variant markers from the Illumina^©^ 550K marker panel. For these variants, we systematically searched for all three-marker syntheses in a 2 Mb surrounding interval in the CEU subsample (85 individuals) of the 1,000 Genomes Project phase 1 integrated release [21] reference data (accessed Mar2012). In table 1 we list the absolute numbers and proportions of syntheses for |*r_xh_*| of either 0.995 or 0.975. A portion of 74.4% (55.8%) of tag variant markers allowed a synthesis by three surrounding variants at an *r_xh_* cutoff of 0.975 (0.995). We investigated how many of these syntheses comprised “broad-sense” functional variants, i.e. variants classified not as ‘unknown’, ‘intergenic’ or ‘intronic’. 55.3% (36.9%) of variants allowed a synthesis including at least one functional variant, 15.7% (8.2%) allowed a synthesis with at least two functional variants and 5.1% (3.4%) could be explained by a synthesis entirely build by functional variants. Thus, formal synthesis, also involving functional variants, in general, is a common phenomenon.

**Table 1:** Number and proportion of syntheses for three-variants sets

| #<sup>a</sup> | $\vert r_{xh}\vert $ | Random | | Crohn’s Disease | | | Type 2 Diabetes | | |
| --- | --- | --- | --- | --- | --- | --- | --- | --- | --- |
|  |  | amount | frequency | amount | frequency | P<sup>b</sup> | amount | frequency | P<sup>b</sup> |
| 3 | 0.995 | 34 | 0.034 | 7 | 0.100 | 5.42e-3 | 5 | 0.083 | 4.86e-2 |
| 3 | 0.975 | 51 | 0.051 | 12 | 0.171 | **3.50e-5** | 5 | 0.083 | 2.77e-1 |
| 2 | 0.995 | 82 | 0.082 | 20 | 0.286 | **2.01e-8** | 13 | 0.217 | **3.90e-4** |
| 2 | 0.975 | 157 | 0.157 | 26 | 0.371 | **4.10e-6** | 16 | 0.267 | 2.55e-2 |
| 1 | 0.995 | 369 | 0.369 | 36 | 0.514 | 1.54e-2 | 27 | 0.450 | 2.08e-1 |
| 1 | 0.975 | 532 | 0.532 | 46 | 0.657 | 4.22e-2 | 40 | 0.667 | 4.20e-2 |
| 0 | 0.995 | 558 | 0.558 | 45 | 0.643 | 1.66e-1 | 37 | 0.617 | 3.74e-1 |
| 0 | 0.975 | 744 | 0.744 | 52 | 0.743 | 9.83e-1 | 48 | 0.800 | 3.32e-1 |
| **total** |  | 1000 |  | 70 |  |  | 60 |  |  |

<sup>a</sup> number of “broad-sense” functionals; <sup>b</sup> compared to random

Next, we investigated the frequency of syntheses for Crohn’s Disease and Type 2 Diabetes susceptibility loci. We took 71 consensus variants for Crohn’s Disease [25] and 62 for Type 2 Diabetes [26] from the GWAS Catalog [27]. Of these variants, 70 Crohn’s variants and 60 Type 2 Diabetes variants were available in the 1,000 Genomes CEU reference data. For those loci our method was able to reveal synthesizing sets, we list the respective rs-numbers, the top set and the correlation coefficient in supplemental table A. The portion of syntheses, ignoring functional annotations, for Crohn’s Disease, 74,3% (64,3%) and Type2 Diabetes, 80.0% (61.7%), did not differ significantly from the portions observed for random variants (*P >* 0.05). However, a substantial increase in the portion of synthetic associations involving functional variants was observed for the susceptibility loci of both phenotypes. After adjustment for multiple comparisons, four settings showed significance: 17.1% of the Crohn’s Disease variants allowed a synthesis with three functional variants at *r_xh_* = 0.975 as compared to a portion of 5.1% for random variants (*P* = 3.5 · 10^−5^). 37.1% (28.6%) of Crohn’s Disease variants allowed a synthesis with at least two functional variants at *r_xh_* = 0.975 (0.995) which reflects a strongly significant increase as compared to random variants (*P* = 2.01 · 10^−8^ and *P* = 4.1 · 10^−6^). Finally, Type 2 Diabetes showed a portion of 21.7% synthesis involving at least two functional variants at *r_xh_* = 0.995 (*P* = 3.9 · 10^−4^). The nominally highly significant cells for both disease phenotypes are highlighted in table 1.

In summary, table 1 shows a significant increase of syntheses involving functional variants for Crohn’s Disease and Type 2 Diabetes susceptibility loci which suggests that a portion of these syntheses potentially reflects the actual functional causes behind the respective GWAS association signal.

### 3.2. Examples of associated susceptibility loci for Crohn’s Disease and Type 2 Diabetes

We further restricted the syntheses discovered in section 3.1 for “narrow-sense” functional variants, i.e. ‘missense’, ‘nonsense’, ‘stop-loss’, ‘frameshift’ and ‘splice-site’ variants. We discovered several complete or nearly complete syntheses made up from variants of those annotation types. In the following we will describe three examples in detail.

First, rs11564258 is an intron variant located in L0C105369736/MUC19 on chromosome 12 which has previously been identified as a susceptibility locus for Crohn’s Disease [25, 27]. Its minor A-allele conveys an odds ratio of 1.73 [1.55;1.95] [25]. Synthesis analysis revealed twelve different three-variant functional syntheses for rs11564258 with |*r_xh_*| ≥ 0.99. A list of these sets can be found in supplemental table B. The synthesizing sets partially overlapped and were made up by a total of 14 different missense variants located in MUC19. Parts of these variants have already been reported in [14]. In order to disentangle the complex LD pattern, we determined the joint haplotype distribution of the tag variant and the synthesizing variants. The respective 15-variant haplotypes with a total length of 138 kb can be found in table 3 and the description of the variants in table 2. The A-allele of the tag variant perfectly tags a single haplotype of frequency 0.024 which fits the previously reported [25] minor allele frequency of 0.025 for rs11564258. While all 14 variants that are involved in the synthesis are missense variants, the tagged haplotype, marked by the red A-allele of the tag variant, does not always carry the minor allele of these variants. Actually, the tagged haplotype carries at most positions the major allele with the exception of rs1444220. Under the convention that the minor allele is the missense allele this suggests that the missense alleles are protective alleles: all further haplotypes carry at least one of these “protective” alleles and the tag haplotype is characterized by an absence of protective alleles. We note, however, that the classification into risk and protective alleles is to a large extent a matter of terminology to describe one of the two sides of a coin. In any case, it can be stated that rs11564258 risk allele carriers can fully be characterized by the allele patterns present at 14 missense variants in MUC19. As shown above, actually various subsets made of three variants are sufficient to obtain the one-to-one correspondence. All of these three-variant-sets contain rs1492319 and rs1444220, while the remaining 12 missense variants are equally well suited to complete the synthesis, see supplemental table B.

**Table 2:** Single variant information

| ID | chr:pos | gene | function | alleles<sup>a</sup> | maf | D<sup>b</sup> | r<sup>2b</sup> |
| --- | --- | --- | --- | --- | --- | --- | --- |
| Synthesizing variants for rs11564258 (LOC105369736/MUC19, intron; Crohn's Disease) |  |  |  |  |  |  |  |
| rs11564258 | 12:40792300 | LOC105369736/MUC19 | intron/intron | A/G | **0.024** | tag variant |  |
| rs4768261 | 12:40834918 | LOC105369736/MUC19 | intron/missense | T/C | 0.171 | -0.70 | 0.00 |
| rs11564109 | 12:40837898 | LOC105369736/MUC19 | missense/upstr 2kb | A/G | rs4768261, r<sup>2</sup> = 1 |  |  |
| rs2933353 | 12:40857943 | MUC19 | missense | A/C | 0.259 | -1.00 | 0.01 |
| rs73100423 | 12:40876068 | MUC19 | missense | A/G | rs4768261, r<sup>2</sup> = 1 |  |  |
| rs2933354 | 12:40876817 | MUC19 | missense | A/G | rs2933353, r<sup>2</sup> = 1 |  |  |
| rs1844979 | 12:40877484 | MUC19 | missense | G/C | rs4768261, r<sup>2</sup> = 1 |  |  |
| rs73100438 | 12:40878665 | MUC19 | missense | A/G | 0.176 | -0.78 | 0.00 |
| rs2588413 | 12:40878869 | MUC19 | missense | C/G | rs2933353, r<sup>2</sup> = 1 |  |  |
| rs2588414 | 12:40879387 | MUC19 | missense | A/G | rs2933353, r<sup>2</sup> = 1 |  |  |
| rs4768276 | 12:40882155 | MUC19 | missense | T/C | rs4768261, r<sup>2</sup> = 1 |  |  |
| rs73114284 | 12:40909924 | MUC19 | missense | T/G | rs4768261, r<sup>2</sup> = 1 |  |  |
| rs1492319 | 12:40918345 | MUC19 | missense | A/G | 0.194 | -1.00 | 0.01 |
| rs1444220 | 12:40922104 | MUC19 | missense | C/A | 0.388 | 1.00 | 0.04 |
| rs1873674 | 12:40930284 | MUC19 | missense | G/C | rs4768261, r<sup>2</sup> = 1 |  |  |
| Synthesizing variants for rs2797685 (PER3, intron; Crohn's Disease) |  |  |  |  |  |  |  |
| rs228682 | 1:7856346 | PER3 | intron | C/T | 0.394 | -1.00 | 0.13 |
| rs34433622 | 1:7861153 | PER3 | intron | C/CT | 0.435 | 1.00 | 0.26 |
| rs7550657 | 1:7876166 | PER3 | intron | T/C | 0.440 | 1.00 | 0.25 |
| rs2797685 | 1:7879063 | PER3 | intron | T/C | **0.165** | tag variant |  |
| rs10462020 | 1:7880683 | PER3 | missense | G/T | 0.165 | -1.00 | 0.04 |
| rs228693 | 1:7882455 | PER3 | intron | A/G | 0.412 | 1.00 | 0.28 |
| rs2859387 | 1:7887248 | PER3 | coding-synon | A/G | 0.392 | 1.00 | 0.31 |
| rs228696 | 1:7887493 | PER3 | missense | T/C | 0.053 | -1.00 | 0.01 |
| rs228697 | 1:7887579 | PER3 | missense | G/C | 0.088 | -1.00 | 0.02 |
| rs2640909 | 1:7890117 | PER3 | missense | C/T | 0.253 | -1.00 | 0.07 |
| rs228665 | 1:7890956 | PER3 | intron | G/C | 0.359 | -1.00 | 0.11 |
| rs4523534 | 1:7894789 | PER3 | intron | G/A | 0.306 | -1.00 | 0.09 |
| rs10462021 | 1:7897133 | PER3 | missense | G/A | rs10462020, r<sup>2</sup> = 1 |  |  |
| rs12029963 | 1:7902231 | PER3 | intron | G/A | 0.312 | -1.00 | 0.09 |
| rs2890565 | 1:7909737 | UTS2 | missense | T/C | 0.029 | -1.00 | 0.01 |
| rs34305100 | 1:7913029 | UTS2 | missense | G/A | 0.212 | -1.00 | 0.05 |
| rs13306061 | 1:7913445 | UTS2 | missense | T/C | 0.212 | -1.00 | 0.05 |
| Synthesizing variants for rs7178572 (HMG20A, intron; Type 2 Diabetes) |  |  |  |  |  |  |  |
| rs1867780 | 15:77407114 | PEAK1 | missense | G/C | 0.306 | -1.00 | 0.21 |
| rs7178572 | 15:77747190 | HMG20A | intron | A/G | **0.324** | tag variant |  |
| rs7119 | 15:77777632 | HMG20A | utr-3' | T/C | 0.371 | -1.00 | 0.28 |
| rs144284088 | 15:78290563 | TBC1D2B | missense | T/C | 0.041 | -1.00 | 0.02 |
| rs2229961 | 15:78880752 | CHRNA5 | missense | A/G | 0.024 | -0.95 | 0.01 |
| rs189146505 | 15:79058730 | ADAMTS7 | missense | G/A | 0.029 | -1.00 | 0.01 |
| rs186123571 | 15:79059335 | ADAMTS7 | missense | A/G | 0.024 | -1.00 | 0.01 |
| rs61752773 | 15:79089074 | ADAMTS7 | missense | T/C | 0.041 | -1.00 | 0.02 |
| rs28603308 | 15:79284127 | RASGRF1 | missense | A/C | 0.029 | -1.00 | 0.01 |

<sup>a</sup> minor/major allele; <sup>b</sup> with the tag variant

**Table 3:**
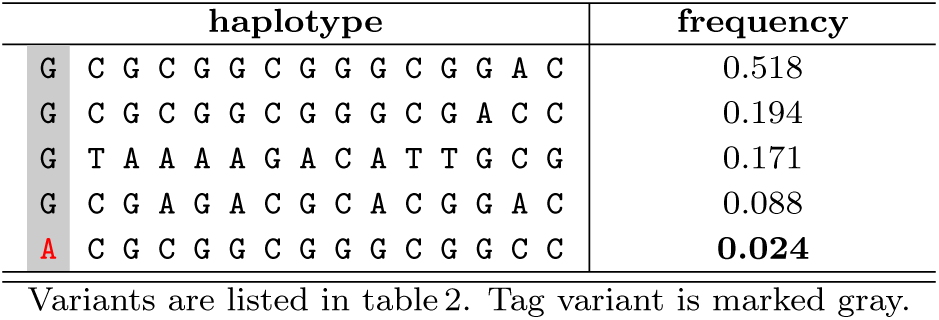
Haplotypes for rs11564258 (LOC105369736/MUC19, intron; Crohn’s Disease) and synthesizing variants

Second, rs2797685 is an intron variant located in PER3 on chromosome 1 which has previously been identified as a susceptibility locus for Crohn’s Disease [25, 27]. Its minor T-allele conveys an odds ratio of 1.05 [1.01-1.10] [25]. Synthesis analysis revealed a manifold of 40 syntheses at |*r_xh_*| ≥ 0.99 with a cardinality between 3 and 6 variants. The synthesizing sets are given in supplemental table B and each of them consists of one non-functional variant in the “narrow-sense” and exclusively additional missense variants. Information of the variants are given in table 2 and the 17-variant haplotypes of the joint distribution with a total length of 56 kb can be found in table 4 (upper panel). The tagged haplotype carries at most major alleles but 4 minor alleles of the variants rs34433622, rs7550657, rs228693 and rs2859387. Since this haplotype comprises a number of intron variants and a synonymous coding variant, we checked if the “narrow-sense” variants are sufficient to characterize the tag variant. This 9-variant haplotype is given in table 4 (lower panel). The yellow marked lines demonstrate a degeneracy of the 8-missense-variant-haplotype which splits up to a *f* = 0.308 haplotype with the C-allele of the tag variant the *f* = 0.165 haplotype comprising the T-allele. There is at least one additional variant necessary, for instance the synonymous coding variant rs2859387, in order to cancel this degeneracy.

**Table 4:**
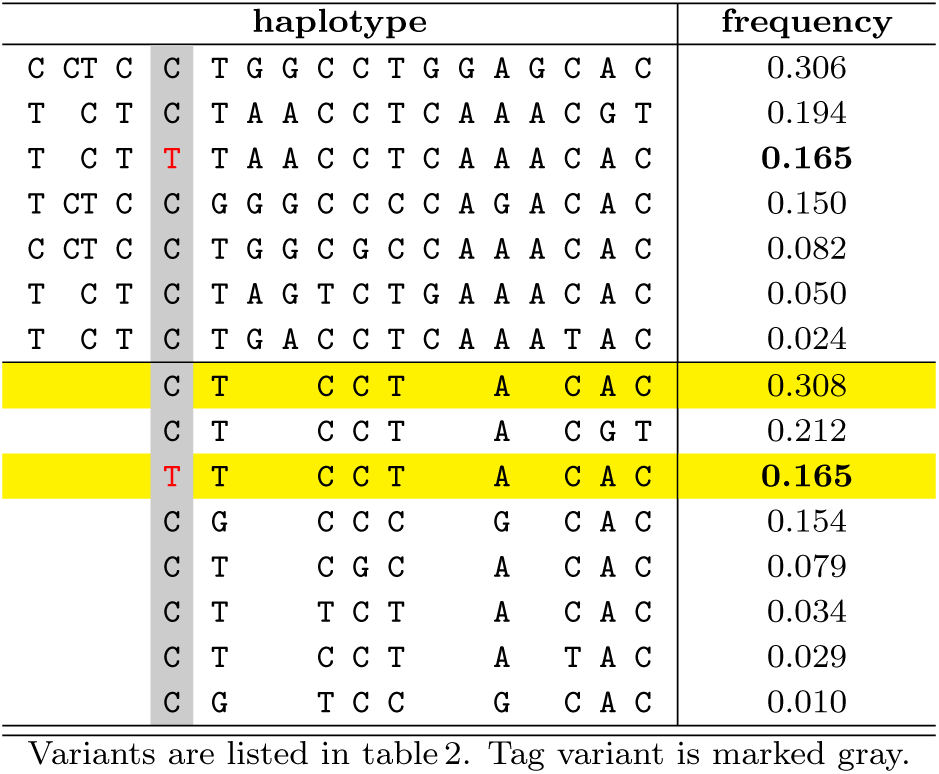
Haplotypes for rs2797685 (PER3, intron; Crohn’s Disease) and synthesizing variants

Third, rs7178572 is an intron variant located in HMG20A on chromosome 15 which has previously been identified as a susceptibility locus for Type 2 Diabetes [26, 27]. Its minor G-allele conveys an odds ratio of 1.08 [1.04-1.13] [26]. Synthesis is established by a two-variant haplotype of size 371 kb comprising the missense variant rs1867780 located in PEAK1 and rs7119 in the untranslated-3’ HMG20A region. In total synthesis analysis revealed 17 synthesizing sets, which are listed in supplemental table B. All sets include the aforementioned two variants and between none and two additional missense variants inside exons of the genes TBC1D2B, CHRNA5, ADAMTS7 and RASGRF1. In table 5 we list the 1,877 kb joint nine-variant haplotypes of the tag and all synthesizing variants. The A-allele of the tag variant perfectly tags a single haplotype of frequency 0.324 consisting of the wildtype alleles of all contributing variants.

**Table 5:** Haplotypes for rs7178572 (HMG20A, intron; Type 2 Diabetes) and synthesizing variants

| haplotype |  |  |  |  |  |  |  |  |  |  |  |  | frequency |
| --- | --- | --- | --- | --- | --- | --- | --- | --- | --- | --- | --- | --- | --- |
| C | A | C | C | G | A | G | C | C |  |  |  |  | **0.324** |
| C | G | T | C | G | A | G | C | C |  |  |  |  | 0.313 |
| G | G | C | C | G | A | G | C | C |  |  |  |  | 0.210 |
| C | G | T | T | G | A | G | C | C |  |  |  |  | 0.025 |
| C | G | T | C | G | A | G | T | C |  |  |  |  | 0.020 |
| G | G | C | C | G | A | G | C | A |  |  |  |  | 0.018 |
| G | G | C | C | G | G | G | C | C |  |  |  |  | 0.018 |
| G | G | C | C | A | A | G | C | C |  |  |  |  | 0.018 |
| G | G | C | C | G | A | G | T | C |  |  |  |  | 0.015 |
| G | G | C | T | G | A | G | C | C |  |  |  |  | 0.010 |

Variants are listed in table2.
Tag variant is marked gray.

## 4. Discussion

We presented methods for the exhaustive search of multi-locus haplotype markers in near perfect LD with a tag variant. Such haplotype markers fulfill the formal criteria of a synthetic association [6]. Our filtering criteria, which we deducted heuristically from typical examples of synthetic association presented in [4], based on marker allele frequencies and LD measure criteria effectively reduce the enormously large space of potentially synthesizing variant sets. The approach can be applied in a case-control setting as well as to public reference genotype data. Via these filtering parameters and our ordering algorithm a quasi-exhaustive search for syntheses is made computationally feasible. Inferring those variants may yield additional insight and help to identify causal genes.

Our data analysis demonstrated that formal synthesis is a very common phenomenon in regions of linkage disequilibrium. We could further show that syntheses involving functional variants occur more frequently with known GWAS susceptibility loci (Crohn’s Disease and Type 2 Diabetes) than with random variants. A potential limitation of this finding is that it is not trivial to select an appropriate set of random variants for comparison. GWAS susceptibility loci are certainly not randomly distributed in the genome. In addition, their allele frequency spectrum will differ from the overall spectrum, alone for the reason of power in discovery studies. In any case, we believe that the detection of syntheses is of substantial relevance, whether or not is possible to proof an enrichment of its frequency among disease associated variants: LD, in general, is a very strong phenomenon and it can be expected that the majority of syntheses found in reference data will carry over to case-control data in the sense that the synthesis will explain the association signal of a tag variant. Verification in case-control data for the phenotype the tag variant is associated with, is, of course, an important validation step. It has to be noted that by statistical means alone it is not possible to ultimately prove causality of a set of variant markers. In the presence of near perfect LD between two or more variant or haplotype markers any of these might provide causality. Still it is important to know all syntheses of a susceptibility locus. First, these syntheses point to functional variants that might underlie the signal. Those variants are at least primary candidates for causality. Second, even if the functional variants of a perfect synthesis are not causal, it has to be acknowledged in functional studies that allele carriers of the tag risk allele also carry the functional variants of the synthesis, and hence are exposed to their biological consequences.

For some analyzed regions we found syntheses involving variants which are as far as 1 Mb or more away from the tag variant. In this sense, we can confirm the statement given in [4] that there might be variants contribution to a synthesis that far away from the main association peak. Of note, the functional synthesis for the diabetes susceptibility locus in HMG20A we described also comprised variants more than 1 Mb away from the tag variant.

In summary, we have shown that the inference of synthetic variants has the potential to yield additional insight into the biology underlying hard-to-interpret association signals. We found a significantly increased number of syntheses involving functional variants for previously confirmed Crohn’s Disease susceptibility loci suggesting that a relevant portion of these reflect the true cause of the association. Furthermore, we detected intriguing examples of syntheses with missense variants for Crohn’s Disease and Type 2 Diabetes. For the future, we plan to apply our approach to all known phenotype associations listed in the GWAS Catalog [27] and to provide a data base of functional syntheses of susceptibility loci. This will display a very valuable resource for the follow-up research of hits from association studies.

Our methods are implemented in the efficient software tool GetSynth, which is freely available and regularly updated at <u>http://sourceforge.net/projects/getsynth/</u>. The software is written in C/C++ and requires the binary genotype files, defined by PLINK [29, 30], as input format. Filter criteria, set size, function classes and the number of functional variants that shall be involved in a synthesis and lots of more option can be pre-specified by the user.

## Supplemental Data

Supplemental Data include two tables.

## Acknowledgements

### Web resources

GetSynth: http://sourceforge.net/projects/getsynth/

1,000 Genomes Project: http://www.1000genomes.org

PLINK2: http://www.cog-genomics.org/plink2/

GWAS CATALOG: http://www.ebi.ac.uk/gwas/

Ensembl: http://www.ensembl.org

## Appendix A. Ordering algorithm

As a first step the data set is pruned for variants with *r*^2^ ≥ *c*, where c has to be chosen *c* ≤ 1. While pruning and setting very narrow allowed LD space in condition 1.) of section 2.1 tapers the number of variants *n* to be tested, a pre-definition of a maximal set size *N* reduces the number of considered sets from the exponential 2*^n^* − 1 to a sum of binomial coefficients 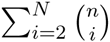 The number of tests to perform are further reduced by condition 2.) and 3.) of section 2.1. Nonetheless the sheer number of possible sets by rare variants from sequencing may become a computational challenge, if the number of provided variants becomes large.

We arrange the list of candidate variants from condition 1.) in decreasing order of |*D*′(*a_i_,a*)|. Our algorithm picks successively one variant at a time and forms all possible sets with all formerly picked variants up to a set cardinality of N. If a set passes the conditions 2.) and 3.) the test is performed. The sets are created in an order, that all subsets have previously been considered. Runtime can further be reduced by additional restrictions: if an annotation file with the genetic functions for the variants is provided, a minimum number of “functional” variants per set can be demanded. Supersets for already detected sets could optionally be omitted.

## Appendix B. Haplotype binary evaluation

We transform the genotypes of a set of variants per individual into binary bitsets storing partial multi-variant genotypes. As a bitset an integer number of sufficient large word length can be used or alternative bitset constructs provided by the employed programming language. First the variants are ordered in an arbitrary way. For missing genotypes we have to omit the individual for the calculation or have to impute missing genotypes in the preface. We use one bitset storing the heterozygous genotypes ‘h1’ and one bitset storing the homozygous two-mutation genotypes ‘h2’. The bitset for the two-wildtype homozygous genotypes ‘h0’ is not needed for the following consideration, but could be obtained by the binary operation NOT(h1 OR h2). An example for these genotypes on eight variants is:

| genotypes |  | bitset |  |  |  |  |  |  |  |
| --- | --- | --- | --- | --- | --- | --- | --- | --- | --- |
| homozygous | 'h0' | 1 | 0 | 0 | 0 | 0 | 1 | 0 | 0 |
| heterozygous | 'h1' | 0 | 0 | 1 | 1 | 0 | 0 | 1 | 0 |
| homozygous | 'h2' | 0 | 1 | 0 | 0 | 1 | 0 | 0 | 1 |

The multi-variant genotype ‘h2’ trivially contributes to both resulting haplotypes in a haplotype pair, but the ‘h1’ part needs to be distributed in any combination (2^checksum^ in total) to one of both haplotypes. In order to avoid idling cycles one should omit considering zeros in ‘h1’ by making use of a map-array. Let now TYPE be any sub-bitset of ones from ‘h1’ then kmpl = h1 XOR type is the complement sub-bitset to type from ‘h1’. The pairs of haplotypes for an individual are simply constructed by h2 OR type and h2 OR kmpl. With all combinations of possible haplotype pairs per individual we can perform the maximum-likelihood estimation of the haplotype frequencies [23, 24]. The expectation maximization recursion formula for the frequencies is given by

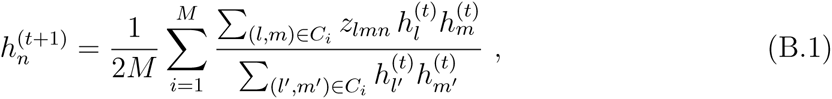

where *M* is the number of individuals, *C_i_* is the set of all ordered pairs of haplotypes compatible with ‘h1’ and ‘h2’ for individual *i*, *h_j_* denotes the frequency of haplotype *j* and *z_lmn_* is an indicator variable, {0,1, 2}, equal to the number of times the haplotype *n* is present in the pair (*l,m*).

## References

[1] Hirschhorn, J.N., Daly, M.J. (2005). Genome-wide association studies for common deseases and complex traits. Nat. Rev. Genet. 6, 95–108 (doi:10.1038/nrg1521).

[2] Bush, W.S., Moore, J.H. (2012). Chapter 11: Genome-Wide Association Studies. PLoS Comput. Biol. 8, e1002822 (doi:10.1371/journal.pcbi.1002822).

[3] Platt, A., Vilhjálmsson, B.J., Nordborg, M. (2010). Conditions under which genome-wide association studies will be positively misleading. Genetics 186, 1045–1052 (doi:10.1534/genetics.110.121665).

[4] Dickson, S.P., Wang, K., Krantz, I., Hakonarson, H., Goldstein, D.B. (2010). Rare variants create synthetic genome-wide associations. PLoS Biology 8, e1000294 (doi:10.1371/journal.pbio.1000294).

[5] Atwell, S., Huang, Y.S., Vilhjlmsson, B.J., Willems, G., Horton, M., Li, Y., Meng, D., Platt, A., Tarone, A. M., Hu, T.T., et al. (2010). Genome-wide association study of 107 phenotypes in Arabidopsis thaliana inbred lines. Nature 465, 627–631 (doi:10.1038/nature08800).

[6] Goldstein, D.B. (2009). Common genetic variation and human traits. N. Engl. J. Med. 360, 1696–1608 (doi:10.1056/NEJMp0806284).

[7] Edge M.D., Gorroochurn P., Rosenberg N.A. (2013). Windfalls and pitfalls: Applications of population genetics to the search for disease genes. Evol. Med. Public Health 2013, 254–272 (doi:10.1093/emph/eot021).

[8] Wang, K., Dickson, S.P., Stolle, C.A., Krantz, I.D., Goldstein, D.B., Hakonarson, H. (2010). Interpretation of association signals and identification of causal variants from genome-wide association studies. Am. J. Hum. Genet. 86, 730–742 (doi:10.1016/j.ajhg.2010.04.003).

[9] Takeuchi, F., Kobayashi, S., Ogihara, T., Fujioka, A., Kato, N. (2011). Detection of common single nucleotide polymorphisms synthesizing quantitative trait association of rarer causal variants. Genome Res. 21, 1122–1130 (doi:10.1101/gr.115832.110).

[10] Wadelius, M., Chen, L.Y., Eriksson, N., Bumpstead, S., Ghori, J., Wadelius, C., Bentley, D., McGinnis, R., Deloukas, P. (2007). Association of warfarin dose with genes involved in its action and metabolism. Hum. Genet. 121, 23–34 (doi:10.1007/s00439-006-0260-8).

[11] Takeuchi, F., McGinnis, R., Bourgeois, S., Barnes, C., Eriksson, N., Soranzo, N., Whittaker, P., Ranganath, V., Kumanduri, V., McLaren, W., et al. (2009). A genome-wide association study confirms VKORC1, CYP2C9, and CYP4F2 as principal genetic determinants of warfarin dose. PLoS Genet. 5, e1000433 (doi:10.1371/journal.pgen.1000433).

[12] Fellay, J., Thompson, A.J., Ge, D., Gumbs, C.E., Urban, T.J., Shianna, K.V., Little, L.D., Qiu, P., Bertelsen, A.H., Watson, M., et al. (2010). ITPA gene variants protect against anaemia in patients treated for chronic hepatitis C. Nature 464, 405–408 (doi:10.1038/nature08825).

[13] Scherag, A., Jarick, I., Grothe, J., Biebermann, H., Scherag, S., Volckmar, A.L., Vogel, C.I., Greene, B., Hebebrand, J., Hinney, A. (2010). Investigation of a genome wide association signal for obesity: synthetic association and haplotype analyses at the melanocortin 4 receptor gene locus. PLoS One 5, e13967 (doi:10.1371/journal.pone.0013967).

[14] Kumar, V., Mack, D.R., Marcil, V., Israel, D., Krupoves, A., Costea, I., Lambrette, P., Grimard, G., Dongm J., Seidman, E.G., et al. (2013). Genome-wide association study signal at the 12q12 locus for Crohn’s Disease may represent associations with the MUC19 gene. Inflamm Bowel Dis. 19, 1254–1259 (doi:10.1097/MIB.0b013e318281f454v).

[15] Orozco, G., Barrett, J.C., Zeggini, E. (2010). Synthetic associations in the context of genome-wide association scan signals. Hum. Mol. Genet. 19, R137–144 (doi:10.1093/hmg/ddq368).

[16] Anderson, C.A., Soranzo, N., Zeginni, E., Barret, J.C. (2011). Synthetic associations are unlikely to account for many common disease genome-wide association signals. PLoS Biol. 9, e1000580 (doi:10.1371/journal.pbio.1000580).

[17] Goldstein, D.B. (2011). The importance of synthetic associations will only be resolved empirically. PLoS Biol. 9, e1001008 (doi:10.1371/journal.pbio.1001008).

[18] Wray, N.R., Purcell, S.M., Visscher, P.M. (2011). Synthetic associations created by rare variants do not explain most GWAS results. PLoS Biol. 9, e1000579 (doi:10.1371/journal.pbio.1000579).

[19] Chang, D., Keinan, A. (2012). Predicting signatures of ’’synthetic associations” and ’’natural associations” from empirical patterns of human genetic variation. PLoS Comput. Biol. 8, e1002600 (doi:10.1371/journal.pcbi.1002600).

[20] Marian, A.J. (2012). Molecular genetic studies of complex phenotypes. Transl. Res. 159, 64–79 (doi:10.1016/j.trsl.2011.08.001).

[21] 1000 Genomes Project Consortium (2012). An integrated map of genetic variation from 1,092 human genomes. Nature 491, 56–65 (doi:10.1038/nature11632).

[22] Excoffier, L., Slatkin, M. (1995). Maximum-likelihood estimation of molecular haplotype frequencies in a diploid population. Mol. Biol. Evol. 12, 921–927.

[23] Long, J.C., Williams, R.C., Urbanek, M. (1995). An E-M algorithm and testing strategy for multiple-locus haplotypes. Am. J. Hum. Genet. 56, 799–810.

[24] Becker, T., Knapp, M. (2004) Maximum-likelihood estimation of haplotype frequencies in nuclear families. Genet. Epidemiol. 27, 21–32.

[25] Franke, A., McGovern, D.P., Barrett, J.C., Wang, K., Radford-Smith, G.L., Ahmad, T., Lees, C.W., Balschun, T., Lee, J., Roberts, R., et al. (2010). Genome-wide meta-analysis increases to 71 the number of confirmed Crohn’s Disease susceptibility loci. Nat. Genet. 42, 1118–1125 (doi:10.1038/ng.717).

[26] DIAGRAM Consortium, AGEN-T2D Consortium, SAT2D Consortium, MAT2D Consortium, T2D-GENES Consortium, Mahajan, A., Go, M.J., Zhang, W., Below, J.E., Gaulton, K.J., et al. (2014). Genome-wide trans-ancestry meta-analysis provides insight into the genetic architecture of type 2 diabetes susceptibility. Nat. Genet. 46, 234–244 (doi:10.1038/ng.2897).

[27] Fernndez-Surez, X.M., Rigden, D.J., and Galperin, M.Y. (2014). The 2014 Nucleic Acids Research Database Issue and an updated NAR online Molecular Biology Database Collection Nucleic Acids Res. 42, D1–D6 (doi:10.1093/nar/gkt1282).

[28] Cunningham, F., Amode, M.R., Barrell, D., Beal, K., Billis, K., Brent, S., Carvalho-Silva, D., Clapham, P., Coates, G., et al. (2015). Ensembl 2015. Nucl. Acids Res. 43 (D1), D662–D669 (doi:10.1093/nar/gku1010).

[29] Purcell, S., Neale, B., Todd-Brown, K., Thomas, L., Ferreira, M.A., Bender, D., Maller, J., Sklar, P., de Bakker, P.I., Daly, M.J., and Sham, P.C. (2007). PLINK: a tool set for whole-genome association and population-based linkage analyses. Am. J. Hum. Genet. 81, 559–575 (doi:10.1086/519795).

[30] Chang, C.C., Chow, C.C., Tellier, L.C.A.M., Vattikuti, S., Purcell, S.M., Lee, J.J. (2015). Second-generation PLINK: rising to the challenge of larger and richer datasets. GigaScience 4, 7 (doi:10.1186/s13742-015-0047-8).

