## Supplemental tables for "Haplotype synthesis analysis in public reference data reveals functional variants underlying known genome-wide associated susceptibility loci"

Supplemental table A: Synthesizing sets for known susceptibility loci for Crohn's Disease and Type 2 Diabetes

| ID | #[a] | synthesizing set | haplotype | $r_{xh}$ |
|---|---|---|---|---|
| Susceptibility loci for Crohn's Disease | | | | |
| rs11167764 | 3/3 | rs75044351, rs449454, rs71583629 | CAA | -0.966 |
| rs12521868 | 3/3 | rs274548, rs2631370, rs3761660 | CTA | -1.000 |
| rs1799964 | 3/3 | rs1800683, rs746868, rs3093540 | GGG | -1.000 |
| rs181359 | 3/3 | rs730882, rs861855, rs861857 | CTG | -0.953 |
| rs12242110 | 3/3 | rs12255327, rs17500653, rs78655239 | GAG | -0.999 |
| rs1998598 | 3/3 | rs12081207, rs1775447, rs4915574 | TTA | -1.000 |
| rs2058660 | 3/3 | rs12081207, rs1775447, rs4915574 | TTA | -1.000 |
| rs2797685 | 3/3 | rs228680, rs4908699, rs228669 | TGC | -1.000 |
| rs3197999 | 3/3 | rs4768, rs35129566, rs6446296 | ATA | -0.969 |
| rs3792109 | 3/3 | rs1816753, rs72976353, rs12476635 | CTT | -0.960 |
| rs780093 | 3/3 | rs1919128, rs780090, rs33994928 | ACC | -0.952 |
| rs8005161 | 3/3 | rs8007437, rs12878879, rs10139328 | AAC | -1.000 |
| rs102275 | 3/2 | rs61896141, rs116980792, rs174570 | AGC | -0.977 |
| rs11871801 | 3/2 | rs8064765, rs1032072, rs11651671 | ACG | -0.970 |
| rs1893217 | 3/2 | rs2847259, rs11080603, rs116860247 | AAA | -1.000 |
| rs3024505 | 3/2 | rs3024509, rs1800893, rs3024498 | ATT | -1.000 |
| rs3764147 | 3/2 | rs10507522, rs9533685, rs41288329 | AGT | -1.000 |
| rs7423615 | 3/2 | rs7581442, rs2114589, rs116493520 | GAA | -0.957 |
| rs10495903 | 3/2 | rs17031056, rs881477, rs116487096 | CCT | -1.000 |
| rs12720356 | 3/2 | rs60542850, rs2304256, rs439025 | GAT | -1.000 |
| rs2076756 | 3/2 | rs1077861, rs3135499, rs75799357 | TAG | -1.000 |
| rs2413583 | 3/2 | rs1047326, rs11089938, rs1569498 | CCC | -0.953 |
| rs3091315 | 3/2 | rs3091318, rs12948058, rs2857657 | TGC | -1.000 |
| rs4409764 | 3/2 | rs888208, rs7899176, rs117462291 | ATT | -1.000 |
| rs694739 | 3/2 | rs7122759, rs1662185, rs544641 | CGG | -1.000 |
| rs7714584 | 3/2 | rs75756583, rs4958846, rs115783200 | GCG | -1.000 |
| Susceptibility loci for Type 2 Diabetes | | | | |
| rs17106184 | 3/2 | rs78012880, rs1278528, rs113601732 | GGG | -1.000 |
| rs17168486 | 3/2 | rs994368, rs10261580, rs11979624 | TGC | -0.998 |
| rs2028299 | 3/3 | rs4932254, rs56171612, rs80068698 | GGT | -1.000 |
| rs2796441 | 3/2 | rs7021934, rs2488274, rs62576224 | ACG | -0.975 |
| rs319598 | 3/2 | rs58488160, rs319599, rs72787107 | GCG | -0.987 |
| rs516946 | 3/2 | rs28411393, rs2439826, rs75802084 | CAC | -1.000 |
| rs7178572 | 3/2 | rs60863150, rs11632729, rs10519162 | CGG | -1.000 |
| rs7612463 | 2/2 | rs6765801, rs112911738 | CC | -1.000 |
| rs7845219 | 3/3 | rs34293080, rs2904889, rs2678832 | TTC | -1.000 |
| rs10507349 | 3/2 | rs12873876, rs1830, rs301047 | GAA | -1.000 |
| rs10510110 | 3/2 | rs7900012, rs10082476, rs4751891 | CAT | -1.000 |
| rs12899811 | 3/3 | rs876874, rs8037137, rs113563465 | ATG | -0.996 |
| rs1552224 | 3/2 | rs12292949, rs6592483, rs12807696 | TAC | -1.000 |
| rs3802177 | 3/2 | rs2466295, rs1578978, rs113040697 | TAA | -0.966 |
| rs4812829 | 3/3 | rs147258225, rs6031545, rs41283268 | AGC | -1.000 |
| rs5215 | 3/3 | rs12791318, rs4756887, rs36101996 | ATC | -0.993 |

[a] number of variants/number of "broad-sense" functional variants

Supplemental table B: Synthesing sets for rs2797685, rs11564258 and rs7178572

| #[a] | synthesizing set | haplotype | length | freq | $r_{xh}$ |
|---|---|---|---|---|---|
| rs11564258 (LOC105369736/MUC19, intron; Crohn's Disease) | | | | | |
| 3/3 | ts2933354, rs1492319, rs1444220 | GGC | 45287 | 0.0262 | -0.9988 |
| 3/3 | ts2588413, rs1492319, rs1444220 | GGC | 43235 | 0.0262 | -0.9988 |
| 3/3 | ts2933353, rs1492319, rs1444220 | CGC | 64161 | 0.0262 | -0.9988 |
| 3/3 | ts2588414, rs1492319, rs1444220 | GGC | 42717 | 0.0262 | -0.9988 |

| | | | | | |
|---|---|---|---|---|---|
| 3/3 | ts73100438, rs1492319, rs1444220 | GGC | 43439 | 0.0237 | -1.0000 |
| 3/3 | ts11564109, rs1492319, rs1444220 | GGC | 84206 | 0.0235 | -1.0000 |
| 3/3 | ts4768261, rs1492319, rs1444220 | CGC | 87186 | 0.0235 | -1.0000 |
| 3/3 | ts1492319, rs1444220, rs1873674 | GCC | 11939 | 0.0235 | -1.0000 |
| 3/3 | ts73114284, rs1492319, rs1444220 | GGC | 12180 | 0.0235 | -1.0000 |
| 3/3 | ts1844979, rs1492319, rs1444220 | CGC | 44620 | 0.0235 | -1.0000 |
| 3/3 | ts73100423, rs1492319, rs1444220 | GGC | 46036 | 0.0235 | -1.0000 |
| 3/3 | ts4768276, rs1492319, rs1444220 | CGC | 39949 | 0.0235 | -1.0000 |
| rs2797685 (PER3, intron; Crohn's Disease) | | | | | |
| 4/3 | rs7550657, rs228696, rs2890565, rs13306061 | TCCC | 37279 | 0.1681 | -0.9995 |
| 4/3 | rs7550657, rs228696, rs2890565, rs34305100 | TCCA | 36863 | 0.1681 | -0.9995 |
| 4/3 | rs2640909, rs228665, rs2890565, rs13306061 | TCCC | 23328 | 0.1688 | -0.9991 |
| 4/3 | rs2640909, rs228665, rs2890565, rs34305100 | TCCA | 22912 | 0.1688 | -0.9991 |
| 5/4 | rs228697, rs228665, rs10462021, rs2890565, rs13306061 | CCACC | 25866 | 0.1689 | -0.9991 |
| 5/4 | rs10462020, rs228697, rs228665, rs2890565, rs13306061 | TCCCC | 32762 | 0.1689 | -0.9991 |
| 5/4 | rs228697, rs228665, rs10462021, rs2890565, rs34305100 | CCACA | 25450 | 0.1689 | -0.9991 |
| 5/4 | rs10462020, rs228697, rs228665, rs2890565, rs34305100 | TCCCA | 32346 | 0.1689 | -0.9991 |
| 5/4 | rs228682, rs228696, rs10462021, rs2890565, rs13306061 | TCACC | 57099 | 0.1687 | -0.9993 |
| 5/4 | rs228682, rs10462020, rs228696, rs2890565, rs13306061 | TTCCC | 57099 | 0.1687 | -0.9993 |
| 5/4 | rs228682, rs228696, rs10462021, rs2890565, rs34305100 | TCACA | 56683 | 0.1687 | -0.9993 |
| 5/4 | rs228682, rs10462020, rs228696, rs2890565, rs34305100 | TTCCA | 56683 | 0.1687 | -0.9993 |
| 3/2 | rs2859387, rs2890565, rs13306061 | ACC | 26197 | 0.1679 | -0.9998 |
| 3/2 | rs2859387, rs2890565, rs34305100 | ACA | 25781 | 0.1679 | -0.9998 |
| 3/2 | rs228693, rs228696, rs13306061 | ACC | 30990 | 0.1680 | -0.9995 |
| 3/2 | rs228693, rs228696, rs34305100 | ACA | 30574 | 0.1680 | -0.9995 |
| 4/3 | rs34433622, rs228696, rs2890565, rs13306061 | CCCC | 52292 | 0.1709 | -0.9987 |
| 4/3 | rs34433622, rs228696, rs2890565, rs34305100 | CCCA | 51876 | 0.1709 | -0.9987 |
| 5/4 | rs228696, rs2640909, rs4523534, rs2890565, rs13306061 | CTACC | 25952 | 0.1706 | -0.9989 |
| 5/4 | rs228696, rs2640909, rs4523534, rs2890565, rs34305100 | CTACA | 25536 | 0.1706 | -0.9989 |
| 6/5 | rs228696, rs228697, rs4523534, rs10462021, rs2890565, rs13306061 | CCAACC | 25952 | 0.1691 | -0.9991 |
| 6/5 | rs10462020, rs228696, rs228697, rs4523534, rs2890565, rs13306061 | TCCACC | 32762 | 0.1691 | -0.9991 |
| 6/5 | rs228696, rs228697, rs4523534, rs10462021, rs2890565, rs34305100 | CCAACA | 25536 | 0.1691 | -0.9991 |
| 6/5 | rs10462020, rs228696, rs228697, rs4523534, rs2890565, rs34305100 | TCCACA | 32346 | 0.1691 | -0.9991 |
| 5/4 | rs228696, rs2640909, rs12029963, rs2890565, rs13306061 | CTACC | 25952 | 0.1728 | -0.9981 |
| 5/4 | rs228696, rs2640909, rs12029963, rs2890565, rs34305100 | CTACA | 25536 | 0.1728 | -0.9981 |
| 6/5 | rs228696, rs228697, rs10462021, rs12029963, rs2890565, rs13306061 | CCAACC | 25952 | 0.1691 | -0.9988 |
| 6/5 | rs10462020, rs228696, rs228697, rs12029963, rs2890565, rs13306061 | TCCACC | 32762 | 0.1691 | -0.9988 |
| 6/5 | rs228696, rs228697, rs10462021, rs12029963, rs2890565, rs34305100 | CCAACA | 25536 | 0.1691 | -0.9988 |
| 6/5 | rs10462020, rs228696, rs228697, rs12029963, rs2890565, rs34305100 | TCCACA | 32346 | 0.1691 | -0.9988 |
| 5/4 | rs228696, rs2640909, rs228665, rs2890565, rs13306061 | CTCCC | 25952 | 0.1706 | -0.9989 |
| 5/4 | rs228696, rs2640909, rs228665, rs2890565, rs34305100 | CTCCA | 25536 | 0.1706 | -0.9989 |
| 6/5 | rs228696, rs228697, rs228665, rs10462021, rs2890565, rs13306061 | CCCACC | 25952 | 0.1677 | -0.9995 |
| 6/5 | rs10462020, rs228696, rs228697, rs228665, rs2890565, rs13306061 | TCCCCC | 32762 | 0.1677 | -0.9995 |
| 6/5 | rs228696, rs228697, rs228665, rs10462021, rs2890565, rs34305100 | CCCACA | 25536 | 0.1677 | -0.9995 |
| 6/5 | rs10462020, rs228696, rs228697, rs228665, rs2890565, rs34305100 | TCCCCA | 32346 | 0.1677 | -0.9995 |
| 4/3 | rs2859387, rs228696, rs2890565, rs13306061 | ACCC | 26197 | 0.1681 | -0.9995 |
| 4/3 | rs2859387, rs228696, rs2890565, rs34305100 | ACCA | 25781 | 0.1681 | -0.9995 |
| 4/3 | rs228693, rs228696, rs2890565, rs13306061 | ACCC | 30990 | 0.1681 | -0.9995 |
| 4/3 | rs228693, rs228696, rs2890565, rs34305100 | ACCA | 30574 | 0.1681 | -0.9995 |
| rs7178572 (HMG20A, intron; Type 2 Diabetes) | | | | | |
| 2/1 | rs1867780, rs7119 | CC | 370518 | 0.3235 | -1.0000 |
| 3/2 | rs1867780, rs7119, rs186123571 | CCG | 1652221 | 0.3235 | -1.0000 |
| 3/2 | rs1867780, rs7119, rs61752773 | CCC | 1681960 | 0.3235 | -1.0000 |
| 4/3 | rs1867780, rs7119, rs186123571, rs61752773 | CCGC | 1681960 | 0.3235 | -1.0000 |
| 3/2 | rs1867780, rs7119, rs189146505 | CCA | 1651616 | 0.3235 | -1.0000 |
| 4/3 | rs1867780, rs7119, rs189146505, rs186123571 | CCAG | 1652221 | 0.3235 | -1.0000 |
| 3/2 | rs1867780, rs7119, rs28603308 | CCC | 1877013 | 0.3235 | -1.0000 |
| 4/3 | rs1867780, rs7119, rs186123571, rs28603308 | CCGC | 1877013 | 0.3235 | -1.0000 |
| 4/3 | rs1867780, rs7119, rs189146505, rs28603308 | CCAC | 1877013 | 0.3235 | -1.0000 |
| 3/2 | rs1867780, rs7119, rs144284088 | CCC | 883449 | 0.3225 | -0.9998 |
| 4/3 | rs1867780, rs7119, rs144284088, rs186123571 | CCCG | 1652221 | 0.3219 | -0.9996 |
| 3/2 | rs1867780, rs7119, rs2229961 | CCG | 1473638 | 0.3235 | -1.0000 |
| 4/3 | rs1867780, rs7119, rs2229961, rs186123571 | CCGG | 1652221 | 0.3235 | -1.0000 |
| 4/3 | rs1867780, rs7119, rs2229961, rs61752773 | CCGC | 1681960 | 0.3235 | -1.0000 |
| 4/3 | rs1867780, rs7119, rs2229961, rs189146505 | CCGA | 1651616 | 0.3235 | -1.0000 |

| | | | | | |
|---|---|---|---|---|---|
| 4/3 | rs1867780, rs7119, rs2229961, rs28603308 | CCGC | 1877013 | 0.3235 | -1.0000 |
| 4/3 | rs1867780, rs7119, rs144284088, rs2229961 | CCCG | 1473638 | 0.3235 | -1.0000 |

[a] number of variants/number of “narrow-sense” functional variants
